# Prevention of thrombocytopenia and thrombosis in heparin-induced thrombocytopenia (HIT) using deglycosylated KKO: A novel therapeutic?

**DOI:** 10.1101/2022.10.19.512755

**Authors:** Amrita Sarkar, Sanjay Khandelwal, Hyunjun Kim, Yves Gruel, Jerome Rollin, Geoffrey D. Wool, Gowthami M. Arepally, Douglas B. Cines, Lubica Rauova, Mortimer Poncz

## Abstract

Heparin-induced thrombocytopenia (HIT) is characterized by mild thrombocytopenia associated with a highly prothrombotic state due to the development of pathogenic antibodies that recognize human (h) platelet factor 4 (PF4) complexed with various polyanions. While non-heparin anticoagulants and intravenous immunoglobulin (IVIG) are the mainstay of care, bleeding may develop, and risk of new thromboembolic events remain. We had described a mouse IgGκ2b antibody KKO that mimics the sentinel features of pathogenic HIT antibodies, including binding to the same neoepitope on hPF4:polyanion complexes. KKO, like HIT IgGs, activates platelets through FcγRIIA and induces complement activation. We now asked whether Fc-modified KKO can be used as a novel therapeutic to prevent or treat HIT. Using the endoglycosidase EndoS, we created deglycosylated KKO (DGKKO). DGKKO bound to PF4-polyanion complexes, and blocked FcγRIIA-dependent activation of PF4 treated platelets by KKO, 5B9 (another HIT-like monoclonal antibody), and isolated IgGs from HIT patients. DGKKO also decreased complement activation and deposition of C3c on platelets. Injection of DGKKO into “HIT mice” lacking mouse PF4, but transgenic for hPF4 and FcγRIIA, prevented and reversed thrombocytopenia when injected before or after KKO, 5B9 or HIT IgG, respectively, in a microfluidic system. DGKKO reversed antibody-induced thrombus growth in HIT mice. In contrast, DGKKO was ineffective in preventing thrombosis by IgG from a patient with the HIT-related disorder, vaccine-induced immune thrombotic thrombocytopenia. Thus, DGKKO may represent a new class of therapeutics for targeted treatment of patients with HIT.

**Key Points:**

- Deglycosylated (DG) KKO can reverse thrombocytopenia in a HIT murine model.
- DGKKO can prevent/reverse thrombosis *in vitro* and in a HIT murine model.

## Introduction

Heparin-induced thrombocytopenia (HIT) is an autoimmune disorder^1^ mediated by platelet activating antibodies that recognized the chemokine, platelet factor 4 (PF4, CXCL4), complexed to heparin or other polyanions^2,3^. Patients with HIT generally develop only mild thrombocytopenia but an unusually intense hypercoagulable state, leading to venous and arterial thrombosis^4^. Present-day therapeutics for HIT center on non-heparin anticoagulants^5^, which carry a risk of untoward bleeding, yet with limited efficacy at preventing new thrombi^6^.

KKO is a HIT-like monoclonal antibody that selectively binds to PF4 complexed to polyanions, such as heparin^7^, and binds to the same epitope as the majority of patient HIT immunoglobulin G (IgG)^8^. KKO activates human platelets, and neutrophils via both FcγRIIA^9,10^ and complement^11,12^ Infusing KKO or HIT IgGs into transgenic mice that express both FcγRIIA and hPF4 recapitulates both the thrombocytopenia and prothrombotic state^13,14^ that characterize the human disease.

We postulated that if KKO’s FcγRIIA and/or complement activation properties were removed by Fc deglycosylation^15^, the modified antibody may be therapeutic by competitively blocking the known HIT antigenic site on PF4 within PF4:polyanion complexes. Since KKO is an IgG_2bκ_ monoclonal antibody^7^, removal of the Fc domain by pepsin-digestion proved not to be feasible^16^. To address this challenge, we modified KKO by deglycosylating its Fc domain at N297, to create deglycosylated KKO (DGKKO) with reduced FcγRIIA and complement activation capacities. We now show that abrogation of Fc-mediated signaling through FcγRIIA and reduction in complement activation by this HIT-like monoclonal antibody not only prevents *in vitro* platelet activation and thrombosis in a photochemically-injured, endothelial-lined microfluidic device, but prevents and even reverses the thrombocytopenia and thrombosis in a murine model of HIT induced by KKO, 5B9 (another HIT-like monoclonal antibody^17^) or IgG isolated from patients with HIT. We also tested whether DGKKO can block thrombosis in a microfluidic system caused by IgGs isolated from a patient with vaccine-induced immune thrombotic thrombocytopenia (VITT), a post-SARSCov2 adenoviral vaccine-induced prothrombotic disorder that also involves antibodies directed at PF4^18^. Our data show that DGKKO is disease-specific in its efficacy.

## Material and Methods

### Human donor samples and mice studied

Human blood (10 ml) was drawn from healthy donor for *in vitro* studies by gravity using a 19-gauge butterfly into 0.38% final concentration sodium citrate from a superficial, upper-extremity vein. The blood sample were stored at RT and were used within 30-60 minutes of being drawn. Deidentified plasma samples were obtained from patients who had a definite diagnosis of HIT or VITT and underwent therapeutic plasmapheresis. IgG was isolated using protein G agarose (ThermoFisher)

Double-transgenic mice, 6-12 weeks old, expressing both hPF4 and FcγRIIA and lacking mouse PF4 (hPF4^+^/ FcγRIIA^+^/*cxcl4*^-/-^, termed HIT mice, were previously described^19^. There was an equal gender distribution for the thrombocytopenia studies, but only male mice can be studied in the cremaster arteriole laser injury thrombosis model; however, we did not note any prior sex difference in thrombosis in passive immunization HIT murine model using a photochemical injury model^14^.

### Antibodies and other reagents

KKO, a mouse IgG 2bκ anti-hPF4/heparin monoclonal antibody and TRA, an isotype-control monoclonal IgG antibody^7^, were purified from hybridoma supernatants, and deglycosylated using EndoS (Genovis) as described^20^. HIT monoclonal antibody 5B9^17^ was a generous gift from J. Rollin and Y. Gruel from UMR CNRS 7292 and Université François Rabelais, Tours, France. F_(ab′)2_ fragments of the monoclonal anti-mouse CD41 antibody MWReg30 (BD Biosciences) and anti-fibrin 59D8^21^ monoclonal antibody (provided by Hartmut Weiler of the Blood Research Institute Versiti, Milwaukee) were labelled with Alexa Fluor 488 and 647 (ThermoFisher), respectively, as described^20^.

Recombinant human (h) PF4 was expressed in Drosophila Expression System (Invitrogen) S2 cells and purified as described^22^. All proteins were tested for size distribution and purity by gel electrophoresis (sodium dodecyl sulfate-polyacrylamide gel electrophoresis) and were confirmed for low endotoxin using ToxinSensor™ Chromogenic LAL Endotoxin Assay Kit (Genscript). Unfractionated heparin (UFH) used was BD PosiFlush™ (500 USP units/5ml, BD).

### Characterization of DGKKO

Binding of DGKKO to PF4 and complexes of PF4 with unfractionated heparin (UFH) complexes was evaluated by an in-house enzyme-linked immunosorbent assay (ELISA). Briefly, 96-well microtiter plates (Nunc Maxisorp; Nunc International) were coated overnight with 100 μl hPF4 (10 μg/ml) with or without UFH (0.2U/ml), followed by blocking unreactive sites with 1% BSA (bovine serum albumin, Sigma-Aldrich) in phosphate-buffered saline (PBS; Invitrogen). KKO or DGKKO was added in serial dilutions (1-1000 ng/ml) for 1 hour at room temperature. After washing out unbound antibody, horseradish peroxidase–labeled goat anti–mouse IgG (1:10000 dilution; Jackson ImmunoResearch) was added followed by TMB peroxidase substrate (KPL) as described^7^. Color development was measured at 450nm using a Spectramax 384 PLUS (Molecular Devices), and the results were analyzed using SoftMax PRO software (Molecular Devices).

### Analysis of PF4:UFH:KKO and PF4:DGKKO complexes in solution using dynamic light scattering (DLS)

Complexes were formed by incubating hPF4 (10μg/ml) with either KKO or DGKKO (35 μg/ml) and UFH (0.2U/ml) in Hank’s Balanced Salt Solution (HBSS, Gibco). DLS studies performed on a fixed scattering angle Zetasizer Nano-ZS system (Malvern Instruments Ltd.) in disposable cuvettes as described^3^. The Z-average size distribution (i.e., hydrodynamic diameters) of particles based on volume, were measured in HBSS at 25°C with a light backscattering of 173°. Up to 25 repetitive measurements were made at the indicated timepoints. Data analysis was performed using the Zetasizer software, version 7.03 (Malvern Instruments Ltd).

### Platelet activation studies

Whole blood samples collected in citrate (0.32% final concentration) were incubated in Ca^++^/Mg^++^ HBSS (Gibco) 1/50 (v/v) in the presence of 10 mM of Gly-Pro-Arg-Pro-NH2 peptide (Bachem H-1998), Allophycocyanin (APC)-labeled anti-hCD41, PE-labeled anti-P-selectin (both BD Biosciences), and FITC-labeled anti-C3c (Abcam) with PF4 (10 μg/ml) and either KKO or DGKKO (0-200 μg/ml) for 45 minutes at room temperature. Cellfix (BD Biosciences) was added 1/1 (v/v) for 15 minutes at room temperature, samples were diluted with 2 volumes HBSS, stored at 4C, and analyzed by flow cytometry (CytoFLEX LX Beckman Coulter) within 24 hours. Platelets were gated based on the forward-scatter and CD41 fluorescence parameters, and binding of anti-C3c, and anti–P-selectin antibodies was quantified as geometric mean fluorescent intensity (MFI) using FlowJo 10.8 software.

### In vitro thrombosis studies under flow

Microfluidic studies were done in 48-well BioFlux plates (Fluxion biosciences) with channels coated with near-confluent human umbilical vein endothelial cells (HUVEC, ATCC), were performed as described^19,23^. The HUVECs were injured using an HXP120C light source with a 475-nm excitation and 530-nm emission filter for 20-30 seconds, while the channel was perfused with hematoporphyrin (50 μg/ml final concentration, Sigma-Aldrich) (Figure Supplement (S) 1A). Whole blood collected in citrate (0.32% final concentration), labelled with 2 mM calcein AM green (ThermoFisher) and supplemented with PF4 (25 μg/ml) and either KKO (50 μg/ml), 5B9 (50 μg/ml) or HIT or VITT IgGs (1 mg/ml) (Figure S1B). After 15 minutes, DGKKO or a deglycosylated isotype control IgG (DGTRA, each at 50 μg/ml) was added and the sample infused into the channel at 10 dyne/cm^2^. Images were taken immediately, and at 5, 10, and 15 minutes after perfusion started. Platelet accumulation on the injured endothelium field was captured using a Zeiss Axio Observer Z1 inverted microscope using Montage Fluxion software and analyzed using ImageJ as described^23^. Confluent, uninjured endothelial-lined channels, exposed to blood that had not had hPF4 or KKO added were used for background to be subtracted from injured vessels exposed to blood that had been exposed to the various experimental conditions.

### Effect of DGKKO on thrombocytopenia in the HIT murine model

Thrombocytopenia was induced in the HIT murine model by intraperitoneal (IP) injection of 200 μg/mouse of either KKO or 5B9 or of 1 mg/mouse polyclonal human HIT or VITT IgGs. DGKKO (200 μg/mouse) was then injected IP or intravenously (IV) at indicated intervals. Platelet counts were measured at baseline and then at 3 through 120 hours post-induction of thrombocytopenia.

### Effect of DGKKO on thrombosis in the HIT murine model

Cremaster arteriole laser injury was performed as described^19^. Alexa Fluor 488-labeled anti-mouse CD41 F(ab′)2 fragments and Alexa Fluor 647-labeled anti-fibrin 59D8 were injected via the jugular vein to label platelets and fibrin, respectively, in thrombi induced with an SRS NL100 pulsed nitrogen dye laser (440 nm) in cremaster arterioles (Figure S1C). Brightfield and fluorescence snapshots of each injury were taken 5 minutes after the initial injury. KKO (20 μg/mouse), 5B9 (20 μg/mouse) or human HIT IgGs (100 μg/mouse) were then injected through a jugular vein to induce a HIT prothrombotic state, and the same clots were reimaged 15 minutes later as we described^19^. DGKKO or DGTRA (20 μg/mouse) was injected 20 minutes after the HIT antibodies, and the injuries were monitored over the ensuing 30 minutes. Brightfield and fluorescence snapshots of the same injuries were then taken to compare platelet deposition before and after induction of a HIT-like lesion and to assess the capacity of DGKKO and DGTRA to interfere with thrombus growth. Fluorescence intensity analysis was done using ImageJ 1.53k and expressed as post antibody/post injury fold change.

### Statistical analysis

Differences between 2 groups were compared using a 2-sided Student t test. Differences between more than 2 groups were determined by 2-way analysis of variance (ANOVA) with the Geisser-Greenhouse correction and Dunnett’s multiple comparison test. All analyses were performed on GraphPad Prism 9.0 (GraphPad Software). Differences were considered significant when the p values were ≤ 0.05.

### Study ethics approval

Human blood for platelet studies was collected after informed consent from healthy, aspirin-free volunteers using a 19-gauge butterfly needle in 129 mM sodium citrate (10:1, v/v) under a protocol approved by the Children’s Hospital of Philadelphia (CHOP) Institutional Review Board for studies involving human subjects and were consistent with the Helsinki Principles. Animal procedures were approved by the Institutional Animal Care and Use Committee (IACUC) at CHOP in accordance with NIH guidelines and the Animal Welfare Act.

## Results

### Deglycosylation removes FcγRIIA-mediated KKO activity and reduces complement activation

As we reported previously, EndoS effectively deglycosylates KKO^20^. DGKKO retained its capacity to bind preferentially to PF4:UFH complexes as compared to PF4 alone, in the same manner as KKO (Figure 1A), and formed ultralarge immune complexes similar in size to PF4:UFH:KKO complexes assessed using DLS (Figure 1B). Deglycosylation effectively abrogated KKO ability to to activate platelets in the presence of supplemented PF4 as measured by P-selectin expression (Figure 1C); and partially inhibited complement activation and deposition of C3c on platelets (Figure 1D).

**Figure 1.**
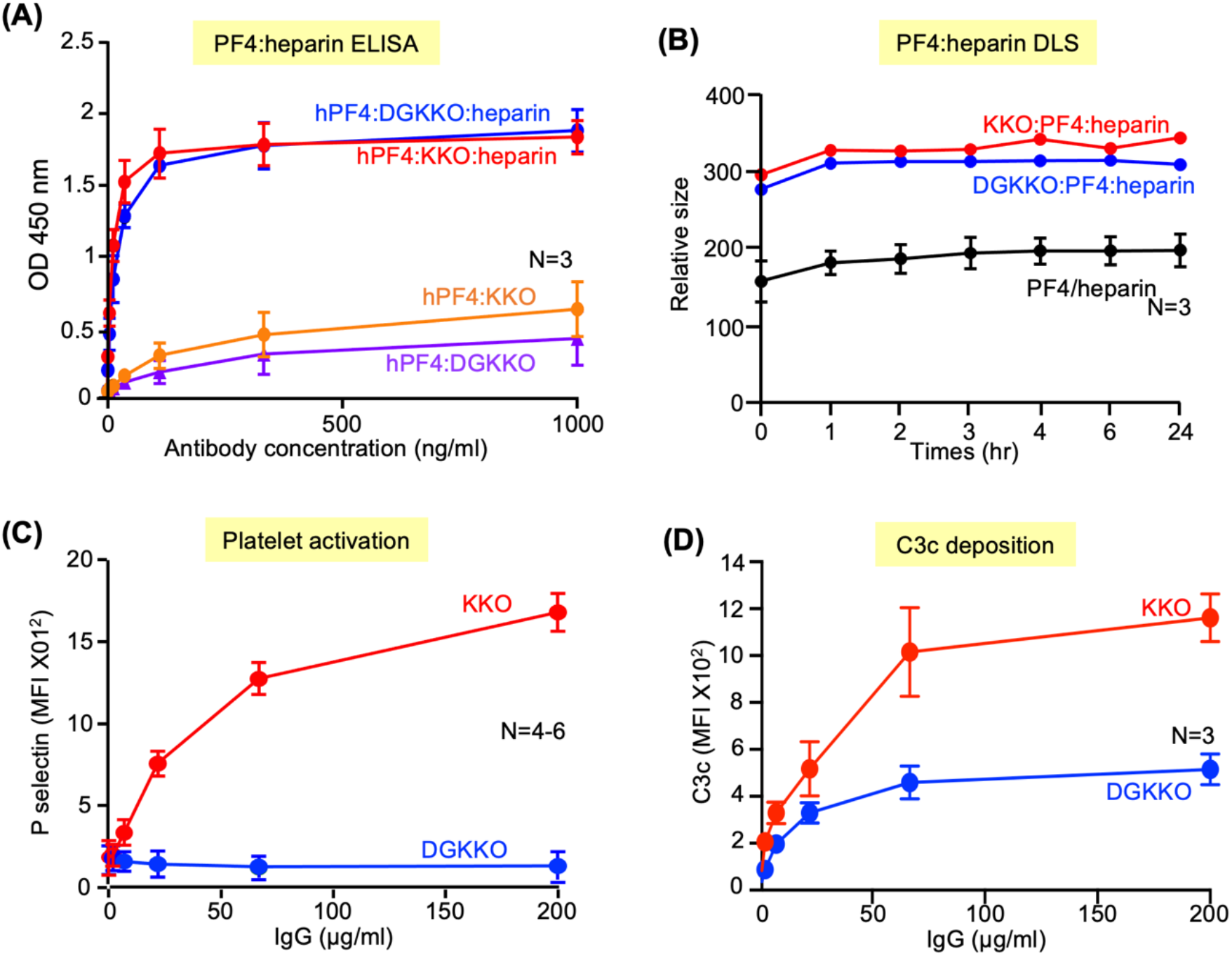
Biological characterization of DGKKO. **(A)** Binding of KKO and DGKKO microtiter wells coated with either hPF4 or hPF4 plus UFH at different antibody concentrations. **(B)** The size of hPF4:UFH:antibody complexes determined by DLS over time in hours (hr). **(C)** Platelet activation KKO vs. DGKKO over a range of antibody concentrations assessed by surface P-selectin expression expressed as Geometric mean of fluorescent intensity (MFI). **(D)** Complement activation as indicated by C3c deposition on the platelet surface in the presence of KKO vs. DGKKO. In **(A)** through **(D)**, the mean ± 1 standard error of the means (SEM) are shown. The number of independent experiments in each study is indicated in the figure. In (C) and (D), p<0.0001 by two-way ANOVA analysis.

### DGKKO prevents thrombosis in a photochemical endothelial injury microfluidic HIT model

To test the ability of DGKKO to prevent thrombosis *in vitro*, we employed an endothelialized microfluidic model previously by us (Figures S1A and S1B)^19,23^. In this model, thrombosis develops when the endothelium had been subject to a photochemical injury and the whole blood samples had been pre-activated by adding hPF4 and HIT antibodies. Adding DGKKO to the human blood 15 minutes post-PF4+KKO activation, markedly reduced platelet accumulation on the injured endothelium (Figures 2A and 2B). Since DGKKO is derived from KKO, we tested another HIT-like monoclonal antibody 5B9^17^ and also polyclonal HIT IgG isolated from plasma of three patients, each of whom met the clinical criteria for HIT, and who had a strongly positive clinical ELISA titer and serotonin-release assay. Similar to KKO, 5B9-induced thrombosis was abrogated by DGKKO. Significant inhibition of platelet adhesion was also seen when thrombosis had been induced by polyclonal HIT IgGs. No significant inhibition of platelet adhesion was seen by isotype control antibody DGTRA (Figure S2A and S2B). On the other hand, DGKKO did not block VITT IgG-mediated thrombosis in the same microfluidic system (Figures 3A and 3B), consistent with the difference in the sites on PF4 recognized by VITT compared with HIT IgG^18^.

**Figure 2.**
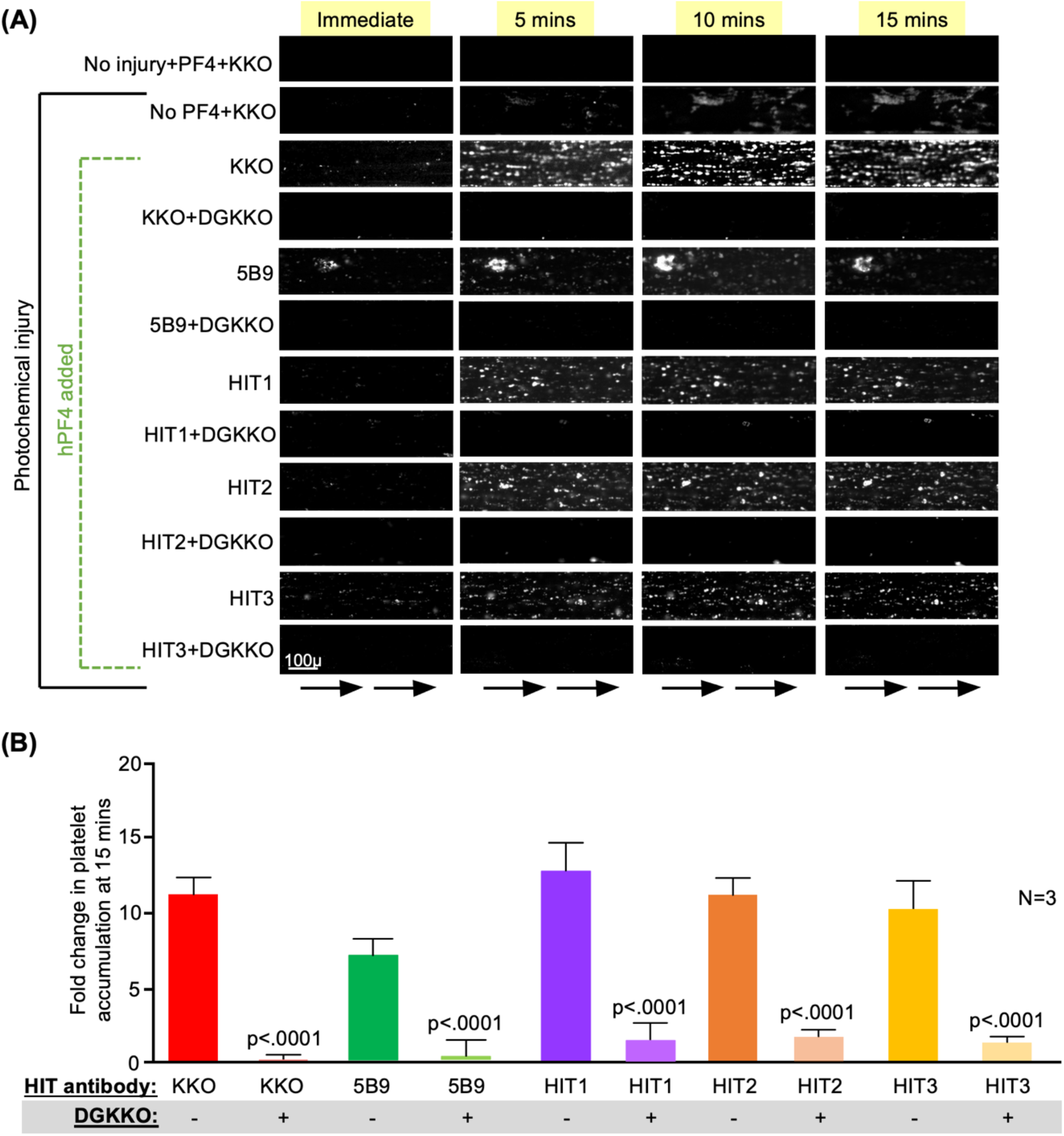
Effect of DGKKO on HIT thrombosis in a microfluidic system. **(A)** Representative images over a 15-minute window of flow through the channels from one of three independent studies showing platelet accumulation on the endothelial lining in white. Size-bar is included and arrows at the bottom indicate direction of flow. Photochemical injury of HUVEC lined channels by hematoporhyrin, supplemental PF4 and HIT monoclonal (KKO and 5B9) or polyclonal (patient derived) antibodies. are necessary to induce platelet adhesion. Effect of DGKKO added 15 minutes after antibody induced activation is shown for each HIT antibody used. **(B)** Quantitative analysis of platelet adhesion showing mean ± 1 SEM of 3 independent experiments. P values were determined by two-way Student t test comparing platelet accumulation in the absence to the presence of DGKKO.

**Figure 3.**
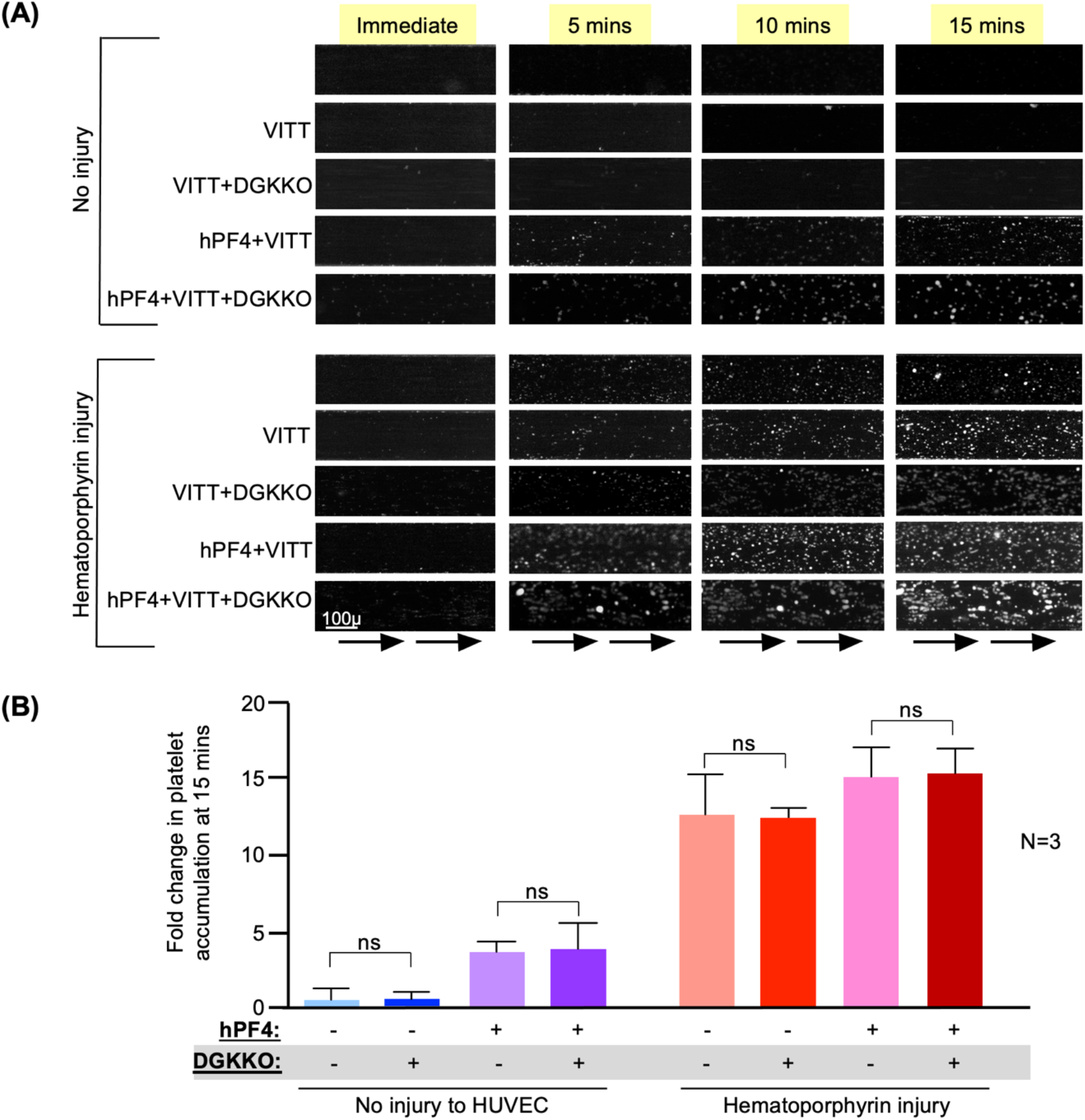
Effect of DGKKO on VITT thrombosis in a microfluidic system. Studies were performed as in Figure 2, but in the presence of IgG isolated from a patient with VITT.

### DGKKO prevents and treats thrombocytopenia in vivo

Thrombocytopenia is a salient feature of HIT. hPF4^+^/FcγRIIA^+^/*cxcl4*^-/-^ (HIT) mice^19^ develop thrombocytopenia by 3 hours after IP injection of KKO or 5B9 or HIT IgG (Figure 4). IP injection of an equal amount of DGKKO 0.5, 1 and 6 hours after IP injection of KKO accelerated platelet recovery (Figure 4A) with maximal effect seen when DGKKO was injected 0.5 hour after KKO. IV injection of DGKKO, 1 hour after IP KKO had been injected, prevented thrombocytopenia from developing and significantly (p<0.05) accelerated platelet recovery. When injected as late as 6 hours after the KKO, accelerated recovery was still seen (Figure 4B). Similar effects by IV DGKKO were seen when thrombocytopenia was induced by 5B9 (Figure 4C) and by polyclonal HIT IgGs (Figures 4D and 4E).

**Figure 4.**
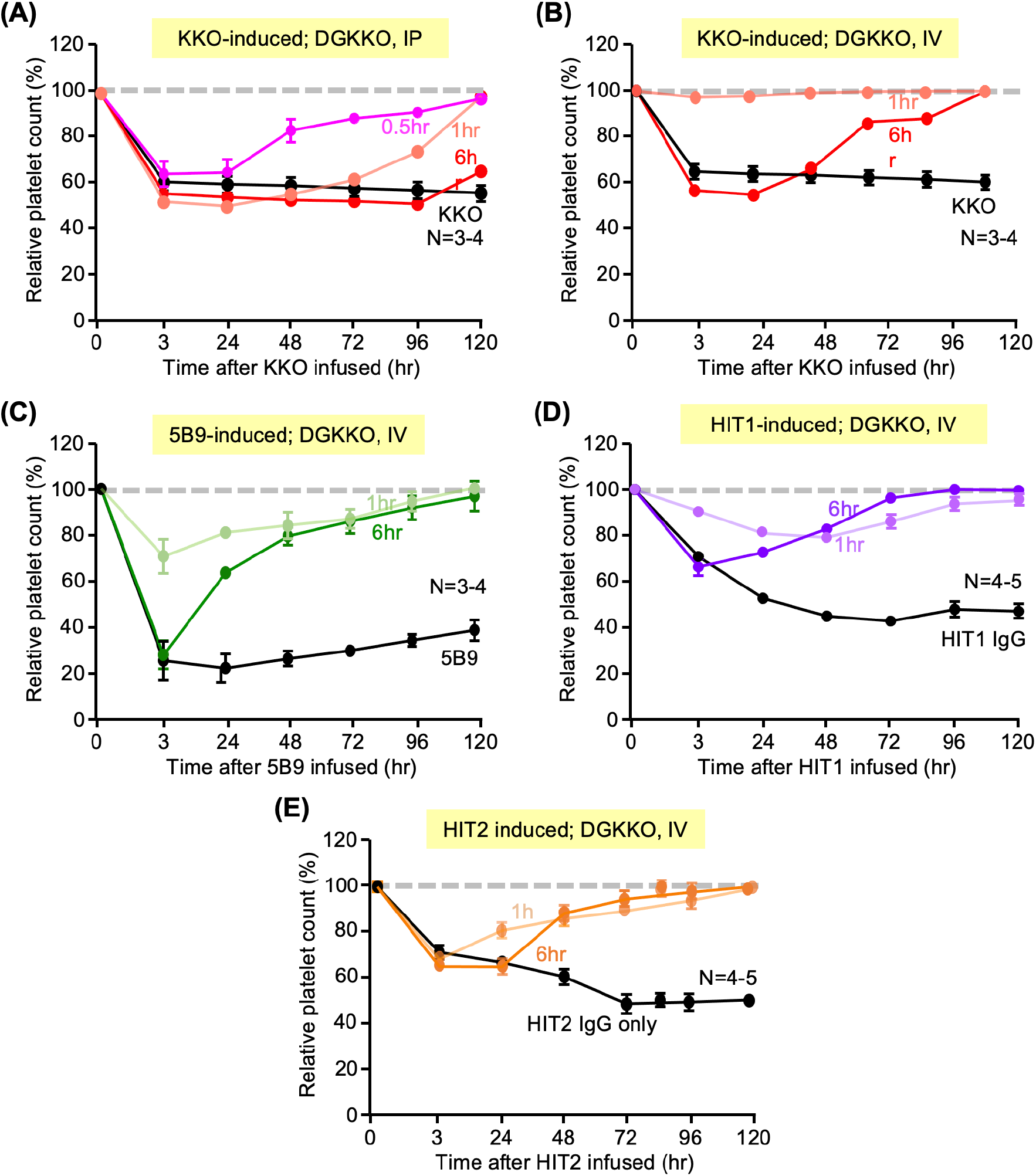
Effects of infused DGKKO in HIT-induced thrombocytopenia in a mice model. Studies were performed in HIT mice after IP injection of HIT-like monoclonal antibody or HIT IgG from each of two patients to induce thrombocytopenia and followed by IP **(A)** or IV **(B)** injection of DGKKO at the indicated times. The number independent experiments for each condition is indicated in the figures. Mean ± 1 SEM of relative platelet counts compared to the baseline are shown. **(A)** Relative platelet counts after IP infusion of KKO followed by IP injection of DGKKO. **(B)** Same as in **(A)**, but DGKKO was given IV. **(C)** Same as in **(B)**, but 5B9 was given instead of KKO. Studies in **(D)** and (**E)** were performed as in **B** using IgG isolated from the plasma of two patients with HIT.

### DGKKO inhibits HIT-induced thrombosis in vivo

HIT is associated with a high risk of limb- or life-threatening thrombosis.^6^ Therefore, we next examined the effect of DGKKO on thrombosis using a cremaster arteriole laser injury model in which we had demonstrated HIT-induced secondary thrombus growth. An initial clot forms at the site of arteriole injury over 5 minutes and then the HIT-like antibody of interest is given IV and results in further thrombus growth, the size of which is measured at 20 minutes after HIT antibodies infusion, just before DGKKO or DGTRA are infused IV (schematic in Figure S1C)^19^. Thrombus size, as indicated by platelet and fibrin accumulation, was measured again after another 30 minutes (Figures 5A). The infusion of DGKKO after KKO caused a reversal in platelet and fibrin accumulation at the site of laser injury (Figure 5A), whereas DGTRA did not (Figure 5B). To determine if the effect of DGKKO was restricted to injury caused by KKO, we tested its effect when 5B9 and a HIT-IgG was infused. These studies showed similar or even greater efficacy in reversing thrombus growth at 30 minutes post-DGKKO infusion based on platelet and fibrin accumulation in both 5B9 and HIT1 IgG studies (Figures 6A and 6B, respectively).

**Figure 5.**
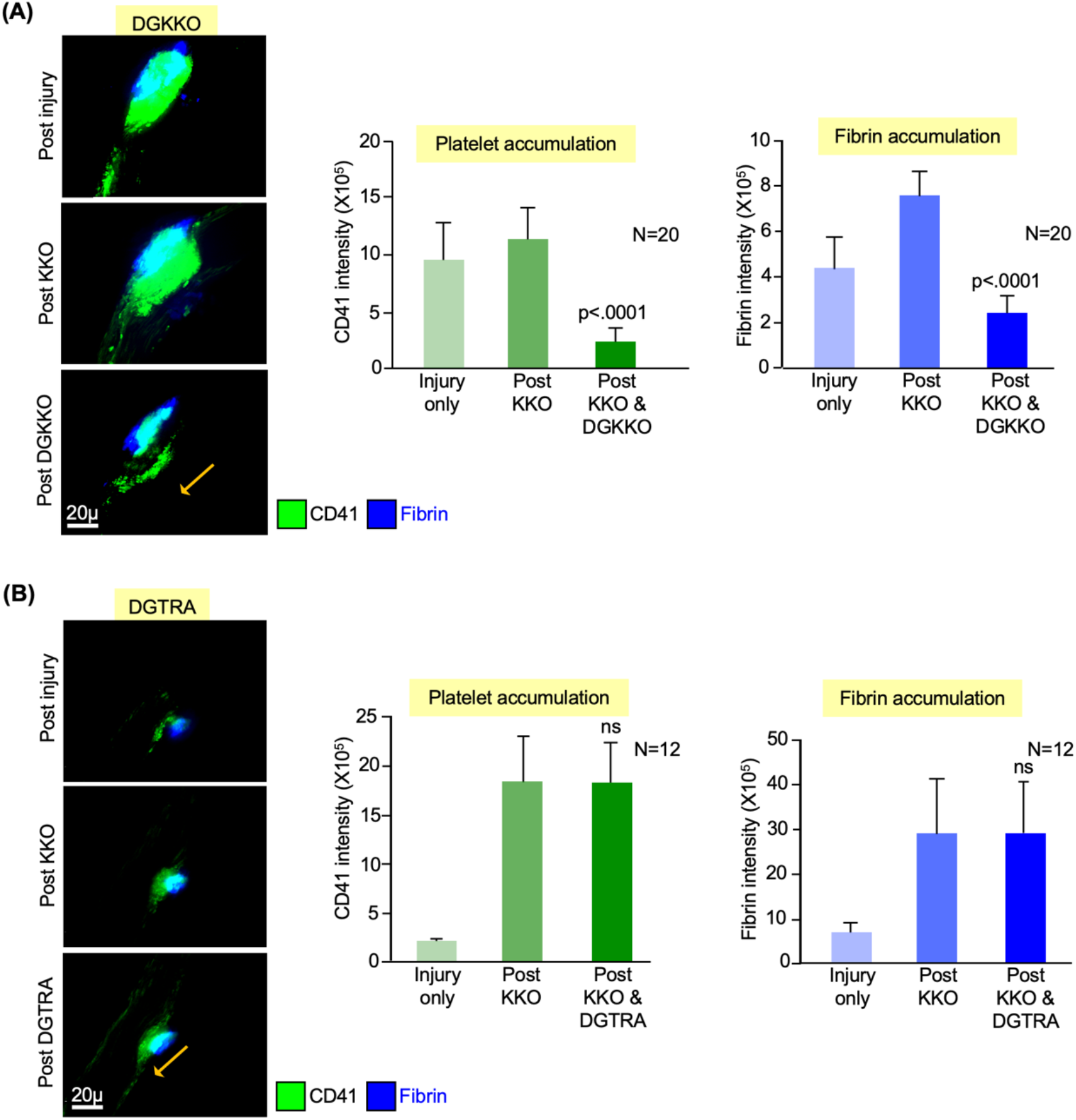
Effect of DGKKO on thrombosis induced by KKO in a murine model of HIT. Cremaster arteriole injury followed by infused KKO HIT prothrombotic studies done as outlined in Supplemental Figure 1C. **(A)** <u>Left</u>: Representative image of 5 mins post-injury vs. 15 mins post-KKO infusion vs. 30 mins post-DGKKO infusion on platelet and fibrin accumulation. Size-bar is included as is arrow indicating direction of flow. <u>Middle</u>: Quantitative analysis of platelet accumulation at the site of injury pre- and post-KKO and post DGKKO. Mean ± 1 SEM are shown. P values were determined by two-way Student t test comparing platelet accumulation in the post-KKO vs. post-DGKKO. <u>Right</u>: Same as middle graph, but for fibrin accumulation. **(B)** Same as **(A)** but DGTRA infused post KKO.

**Figure 6.**
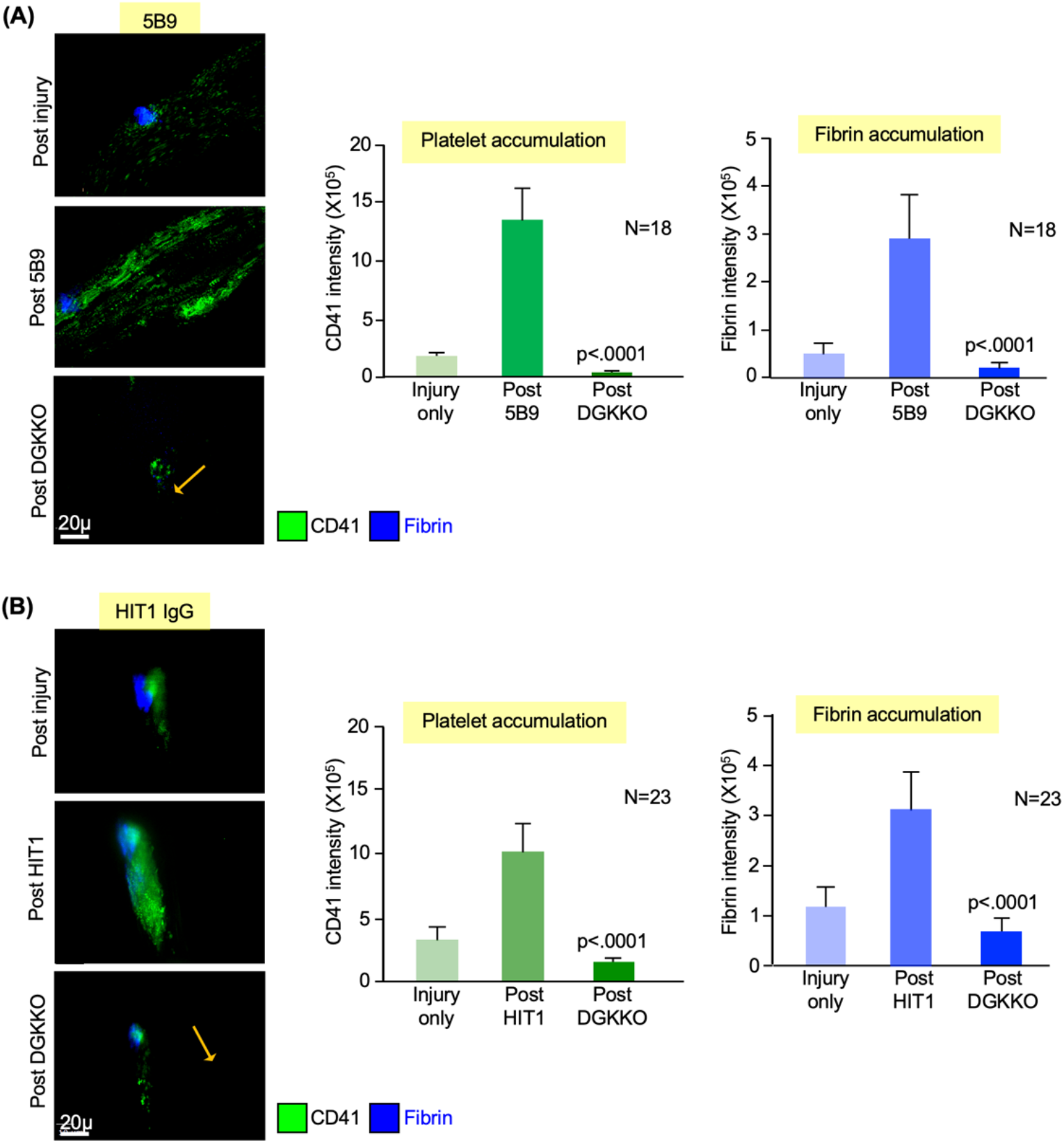
Effect of DGKKO on thrombosis induced by 5B9 and HIT IgG in a murine model of HIT. Thrombus formation in HIT mice was examined as described in the legend to Figure 5 but substituting 5B9 in **(A)** and HIT1 IgG in **(B)**.

## Discussion

Contemporary therapy of HIT is neither entirely effective in terms of preventing new thromboembolic events nor safe in terms of the risk of bleeding^6,24,25^. The critical event for propagation of the intensely prothrombotic state typical in HIT is engagement and crosslinking of Fcγ receptors, predominantly FcγRIIA on platelets and other vascular cells^9,10,26^, by an array of the Fc ends of antibodies that are approximated on ultralarge hPF4:polyanion complexes^27^. The finding that most HIT antibodies and the monoclonal HIT-like antibody KKO compete for binding to PF4^8,28^ led us to the supposition that a KKO antibody analogue that neither binds FcRγIIA, which is required for platelet activation *in vitro*^9^ and the development of thrombosis and thrombocytopenia in murine HIT^13,14^, and that has limited ability to activate complement, which amplifies the autoantibody response^12,29^, might provide a rationale, disease-specific intervention.

We had previously shown that deglycosylation of KKO created a novel monoclonal antibody, DGKKO^20^, that possessed these desirable properties. In this paper we show that deglycosylation of KKO does not affect its binding to hPF4:UFH complexes and maintains the ability to compete with pathogenic human HIT IgG. DGKKO inhibits KKO-induced platelet adhesion to the endothelium under flow in a microfluidic system, and mitigates the severity of thrombocytopenia and prevents thrombotic vascular occlusion in murine HIT. DGKKO also blocked the effector function of 5B9, monoclonal antibody with human IgG1 Fc, shown previously to recognize an epitope that overlaps with KKO and competes with KKO for binding to PF4^17^. Surprisingly, DGKKO, given within 20 min after KKO, actually reversed platelet and fibrin deposition in the microvasculature, suggesting the clots are initially unstable and subject to endogenous dissolution if not organized by accretion of additional platelets and fibrin as a result of unopposed thrombin production. The finding that DGKKO did not prevent clot formation by the one VITT^30^ IgG we studied, is consistent with recent findings that the binding site for VITT and HIT IgG do not overlap^18^. It has been also proposed that the rare cases of autoimmune HIT involve the same antigenic target as in VITT^31,32^, and while antibodies from a larger number of patients need to be tested, it is likely DGKKO would be ineffective in this setting as well. Clinical laboratory studies indicating that the anti-PF4 antibody in a patient is heparin-dependent or competes with KKO, (e.g., using HemosIL, a rapid diagnostic tests for HIT^28^) should be done prior to the initiation of DGKKO or a similar modified HIT-like antibody.

Alternative approaches to mitigating the effect of PF4-anti-PF4 immune complexes have been described. IVIG blocks platelet activation by HIT antibodies and is effective in vivo in at least some patients^33-35^. We do not yet know whether IVIG can reverse thrombosis and thrombocytopenia as we observed with DGKKO. *In vivo* infusion of IdeS, a bacterial protease that cleaves the hinge region of heavy chain IgG, abrogates its ability to bind to FcγR-and Fcγ-dependent cellular activation similarly to DGKKO, prevents thrombus formation and thrombocytopenia in a mouse model induced by 5B9 MoAb^36^, that in contrast to KKO has human IgG1 Fc fragment, sensitive to cleavage. However, the ability of IdeS to block thrombocytopenia or thrombosis induced by patient IgG was not tested. Moreover, IdeS does cleave all IgG and can cause hypogammaglobulinemia^37^ and compromise immune defense against bacterial infections. Moreover, anti-IdeS antibodies are frequent in the general population^38^ and its titer increases after exposure to IdeS^37^. Thus, whether either approaches would be advantageous *in vivo* is not clear at the moment though the IdeS approach may be effective in VITT and autoimmune HIT whereas DGKKO would not.

In summary, a deglycosylated version of the HIT-like monoclonal antibody KKO, DGKKO, appears to be a rational disease-specific intervention because it blocks HIT-IgG binding to platelets and likely to monocytes and neutrophils based on the *in vivo* results reported here. Thrombocytopenia and thrombosis by tested HIT-like monoclonal antibodies and by three HIT IgG preparations could be both prevented prophylactically and reversed by DGKKO. A single VITT IgG tested could not be blocked by DGKKO. Pre-clinical studies of additional HIT IgGs are needed before the therapeutic potential of DGKKO can be estimated.

## Authorship

A.S. carried out and evaluated these studies and prepared the manuscript. S.K. helped in complement studies. H.K. helped in some mice studies. J.R and Y.G. developed 5B9 and edited the manuscript. G.W. provided VITT plasmapheresis sample and edited the manuscript. G.M.A. developed KKO, provided HIT IgG samples, helped in complement study, provided research guidance, and edited the manuscript. D.B.C. provided input in DLS studies, overall research guidance for the manuscript, and edited the manuscript. L.R. helped with characterization of DGKKO, provided research guidance, data interpretation and edited the manuscript. M.P. provided overall conceptual development and direction, data interpretation, and manuscript preparation. None of the authors have any financial or related disclosures to make with respect to this manuscript.

## Acknowledgements

This work was supported by the R01HL151730 (G.M.A, D.B.C. and L.R.) and R35HL150698 (L.R. and M.P.). We would like to thank Drs. Khalil Bdeir, Sergey V Zaytsev and Sergi V. Yarovoi at the Univeristy of Pennsylvania for their help and guidance for the DLS experiments.

