## Supplemental Figures for "Prevention of thrombocytopenia and thrombosis in heparin-induced thrombocytopenia (HIT) using deglycosylated KKO: A novel therapeutic?"

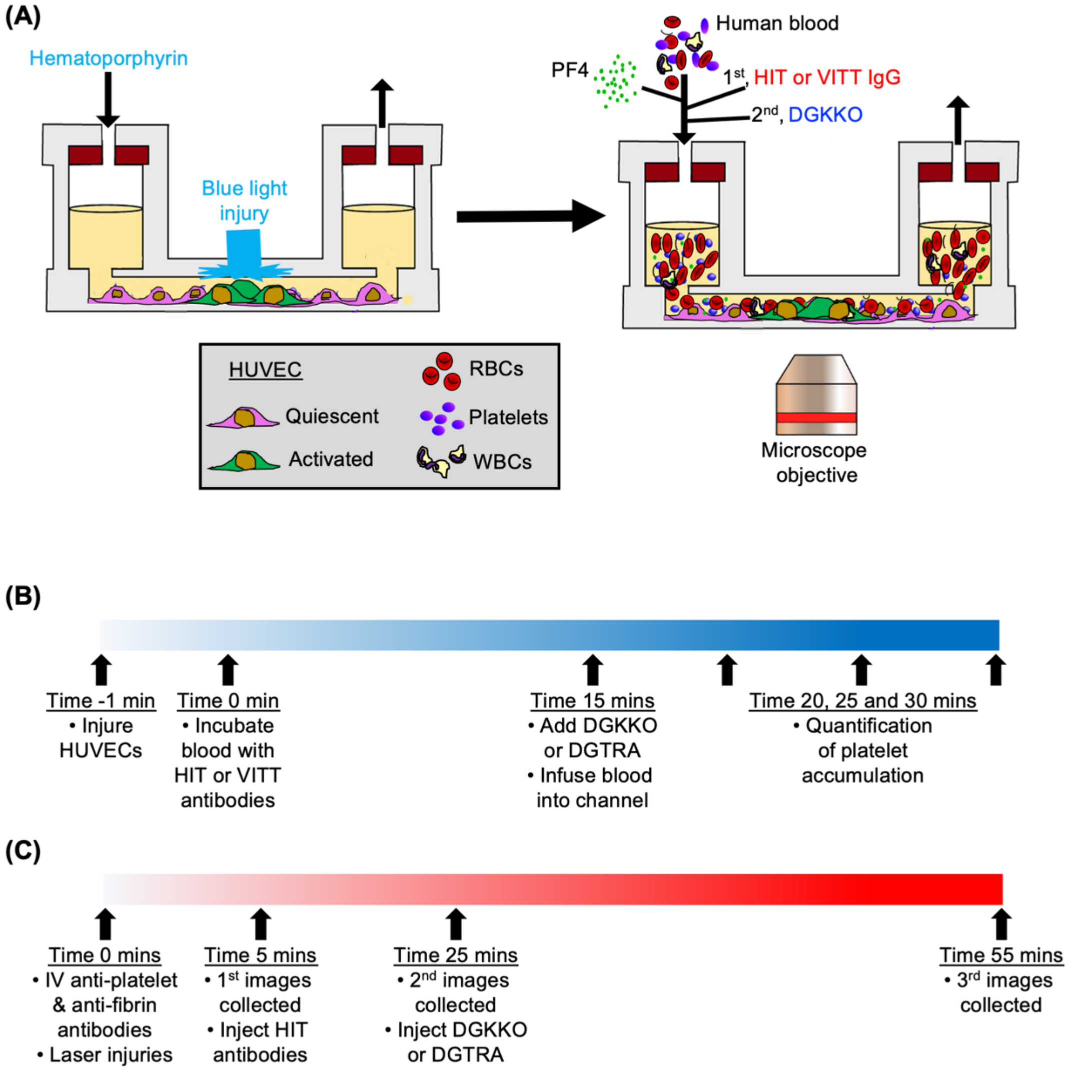


**Figure S1. Schematics of the *in vitro* microfluidic model and the *in vivo* cremaster laser injury models.**

**(A)** and **(B)** are models of the *in vitro* microfluidic system. **(A)** shows the fibronectin-coated, HUVEC-lined system with whole blood flowed through status-post a hematoporphyrin-photochemical injury to part of the channel. Labeled-platelet accumulation during the study is the primary endpoint. **(B)** is the timeline for studies of infused products into the microfluidic channel experiments. **(C)** is the experimental timeline for the procedures and infused products into mouse in the xenotransfusion studies.

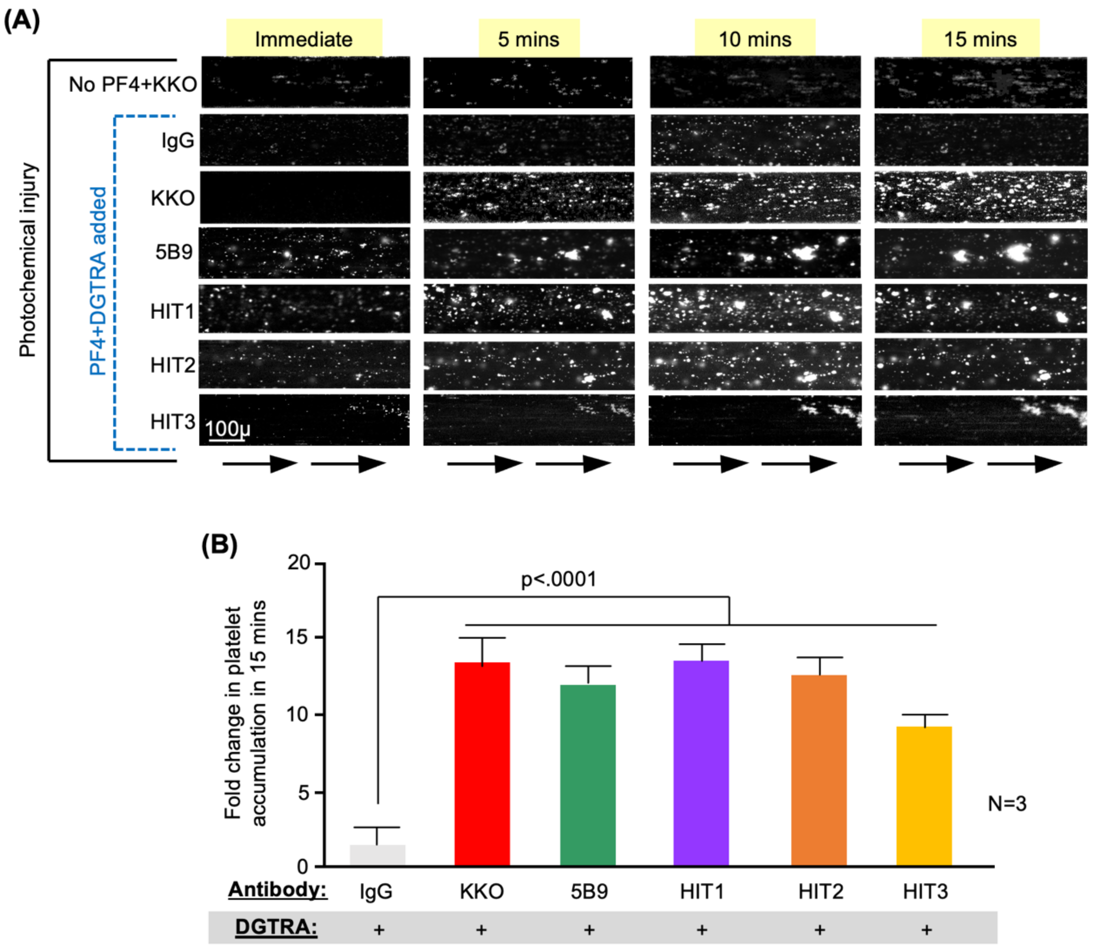


**Figure S2. Effect of DGTRA on HIT thrombosis in a microfluidic system**.

Microfluidic studies as in Figure 2, but with DGTRA isotype control rather than DGKKO. **(A)** Representative images of the channels from one of three independent studies with accumulated platelets on the endothelial lining as white. Size-bar is included and arrows at the bottom indicate direction of flow. **(B)** Summation of studies showing mean ± 1 SEM for the lanes exposed to blood containing DGTRA. P values were determined by two-way Student t test comparing platelet accumulation in the absence to the presence of a HIT-like antibody.
